# Extensile motor activity drives coherent motions in a model of interphase chromatin

**DOI:** 10.1101/319756

**Authors:** David Saintillan, Michael J. Shelley, Alexandra Zidovska

## Abstract

The 3D spatiotemporal organization of the human genome inside the cell nucleus remains a major open question in cellular biology. In the time between two cell divisions, chromatin – the functional form of DNA in cells – fills the nucleus in its uncondensed polymeric form. Recent *in-vivo* imaging experiments reveal that the chromatin moves coherently, having displacements with long-ranged correlations on the scale of microns and lasting for seconds. To elucidate the mechanism(s) behind these motions, we develop a novel coarse-grained active-polymer model where chromatin is represented as a confined flexible chain acted upon by molecular motors, which perform work by exerting dipolar forces on the system. Numerical simulations of this model account for steric and hydrodynamic interactions as well as internal chain mechanics. These demonstrate that coherent motions emerge in systems involving extensile dipoles and are accompanied by large-scale chain reconfigurations and nematic ordering. Comparisons with experiments show good qualitative agreement and support the hypothesis that self-organizing long-ranged hydrodynamic couplings between chromatin-associated active motor proteins are responsible for the observed coherent dynamics.

---

The first real-time mapping of chromatin dynamics across an entire live cell nucleus was recently accomplished using the new method of Displacement Correlation Spectroscopy (DCS) [1]. DCS measurements revealed two types of chromatin motion: a fast local motion that was previously observed by the tracking of single genes [2, 3], and a yet unexplained slower large-scale motion in which parts of the chromatin move coherently across scales of microns and seconds [1]. These coherent motions were found to be activity driven, i.e. dependent upon ATP, but independent of cytoskeletal activity. Furthermore, an analysis of the power spectral density of the chromatin displacement field, under physiological conditions as well as upon ATP-depletion, showed that the active dynamics contributed to chromatin motions at large wavelengths, while short-wavelength fluctuations appeared to be mostly thermally driven [4]. Inhibition of nuclear ATPases (ATP-powered enzymes) such as RNA polymerase II, helicase, and topoisomerase led to the elimination of the coherent motions, suggesting involvement of these molecular motors [1]. The biological implications of these motions are major for the dynamic self-organization of chromatin inside the nucleus and for collective gene dynamics, yet the underlying biophysical mechanisms remain to be revealed.

Motivated by the DCS measurements [1], a first theory of active chromatin hydrodynamics was developed [4]. This theory used a two-fluid model to describe the equilibrium dynamics of the chromatin solution, with chromatin as solute and nucleoplasm as solvent. It introduced two categories of active events: *vector* events describing the effect of force dipoles generated by nuclear enzymes such as polymerases, helicases and topoisomerases, and *scalar* events representing the local condensation and decondensation of chromatin primarily as a result of chromatin remodelers. Scalar events were found to drive longitudinal viscoelastic modes (where the chromatin fiber moves relative to the solvent), while vector events generate transverse modes (where the chromatin fiber moves with the solvent). The theory predicts that chromatin concentration fluctuations dominate at short length scales, while the force-dipole activity dominates the long length scales, implying that the observed coherent motions might indeed be caused by the collective activity of ATP-dependent force-generating nuclear enzymes as supported by experiments [1, 4]. The nature of the force dipoles, i.e., extensile vs contractile, and their ability to drive large-scale reconfigurations of the chromatin fiber via long-ranged nucleoplasmic flows, was not considered.

Active particles that exert extensile force dipoles on a viscous fluid are well known to self-organize through their hydrodynamic interactions and to display strong correlated flows on large length scales [5, 6]. This is especially evident in suspensions of micro-swimmers such as flagellated bacteria which are “pusher” particles [7, 8], and in numerical simulations of many such hydrodynamically interacting swimming cells [9]. The self-organizing effects of extensile force dipoles are also thought to underlie the complex dynamics of cytoplasmic extracts [10, 11] as well as suspensions of *in-vitro* reconstituted microtubules and motor proteins [12, 13]. An additional feature beyond flows is the ordering provided by steric interactions at high concentrations, which leads to liquid-crystalline elasticity and new activity-driven instabilities [14, 15].

In this work, we explore the role of force dipoles in active chromatin dynamics, specifically in driving coherent motions. Since the physical size of active nuclear enzymes such as RNA polymerase II, helicase or topoisomerase is about 5 nm, and the coherent regions span 3 – 5 *μ*m, we hypothesize that micron-scale coherent motions arise from a collective self-organization of active force dipoles acting along and on the chromatin fiber. In contrast to previously studied active matter systems where the force dipoles were free in space (such as swimmers), here the force dipoles are associated to the chromatin fiber. Thus, dipolar self-organization and alignment may result from an interplay of the physical tethering, with the fiber under consequent tensile loads, and of dipolar hydrodynamic interactions mediated by the viscous nucleoplasmic fluid. Further, when packed at high densities, the chromatin fiber may be subject to strong aligning interactions with itself, though these may be frustrated by entanglement. We test these hypotheses using a novel coarse-grained model of the chromatin inside the nucleus as a freely-jointed, spherically-confined chain with transitory force dipoles distributed along its length, and simulate the resulting hydrodynamic flows and chromatin motions.

## EXPERIMENTAL OBSERVATIONS

To gain a mechanistic insight into the nature of coherent chromatin motion, we perform experiments following experimental procedures described in [1] and carry out a new analysis aiming to compare chromatin dynamics in both active (wildtype) and passive (ATP-depleted) states. Using spinning disk confocal microscopy we record chromatin dynamics in HeLa cell nuclei expressing histones H2B-GFP (Fig. 1*A*), and analyze these observations using Displacement Correlation Spectroscopy (DCS) [1]. DCS is a time-resolved image correlation analysis that maps chromatin dynamics simultaneously across an entire live-cell nucleus in real time. It provides maps of local chromatin displacements over time intervals, while sampling all time intervals (and thus time scales) accessible by the experiment (*Materials and Methods*, [1]).

**FIG. 1.**
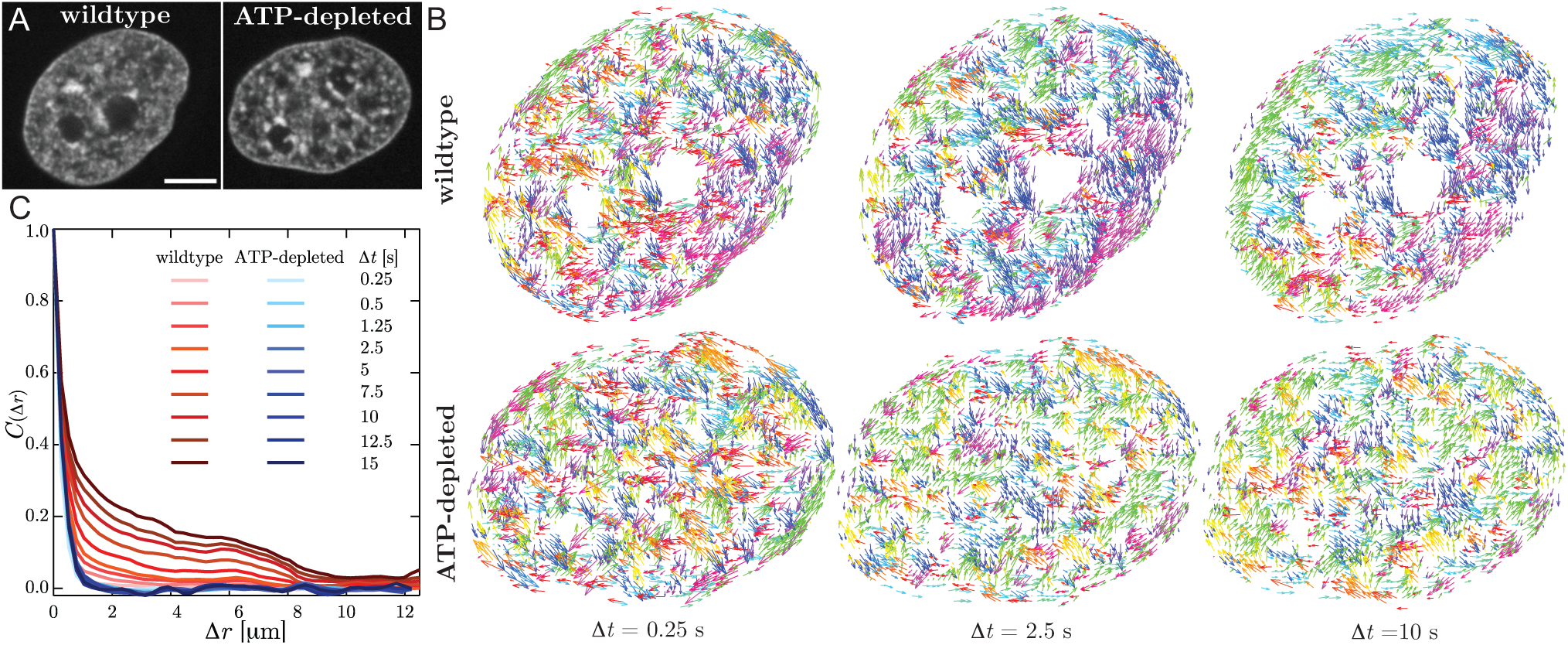
Mapping interphase chromatin dynamics using Displacement Correlation Spectroscopy (DCS) [1]. (*A*) Micrographs of HeLa cell nuclei with fluorescently labeled chromatin (H2B-GFP) for both wildtype and after ATP depletion. (*B*) Displacement fields for the nuclei from *A* for Δ*t* = 0.25, 2.5 and 10 s. Displacement vectors are color-coded by direction. While the wildtype nucleus shows uncorrelated motion at Δ*t* = 0.25 s and an increasingly correlated motion at Δ*t* = 2.5 and 10 s, the ATP-depleted nucleus exhibits uncorrelated motion at all Δ*t*. (*C*) The average spatial displacement autocorrelation functions *C*(Δ*r*) of the DCS maps for the wildtype (*red lines*) and ATP-depleted (*blue lines*) nuclei for different values of Δ*t*. For wildtype, *C*(Δ*r*) shows an increase in spatial correlation with increasing Δ*t*, while upon ATP depletion *C*(Δ*r*) shows a rapid decorrelation at all Δ*t*. Scale bar, 5 *μ*m.

We find that DCS maps for the wildtype and ATP-depleted nuclei are dramatically different (Fig. 1*B*). When we color-code the displacement vectors by their direction, the wildtype nucleus (Fig. 1*B*, *top row*) exhibits displacements spatially uncorrelated in their direction at short time intervals (Δ*t* = 0.25 s), while on longer time scales (Δ*t* = 2.5 and 10 s) patches of correlated displacement vectors emerge (i.e. having the same color) with patch size increasing with Δt. In stark contrast, the ATP-depleted nucleus (Fig. 1*B*, *bottom row*) shows uncorrelated displacements at all time intervals (Δ*t* = 0.25, 2.5 and 10 s). To quantify this observation, we calculate in Fig. 1*C* the spatial displacement autocorrelation function *C*(Δ*r*) of DCS maps for Δ*t* ranging from 0.25 s to 15 s for both wildtype (*red lines*) and ATP-depleted (*blue lines*) nuclei. The difference is striking: for wildtype, *C*(Δ*r*) displays an increasing spatial correlation with increasing Δt, whereas in the ATP-depleted case *C*(Δ*r*) remains unchanged. This implies that ATP-dependent activity drives the coherent motions of chromatin.

These experimental observations motivate us to design a model accounting for the activity in chromatin dynamics, which aims to capture the phenomenology of the experiments. Both the DCS maps as well as *C*(Δ*r*, Δ*t*) allow us to build a direct comparison between the experimental and computational results for both active and passive chromatin dynamics.

## MODEL AND SIMULATION METHOD

We model interphase chromatin as a long flexible Brownian polymer chain acted upon by stochastically-activated dipolar forces. The chain is composed of *N* beads with positions **r**_*i*_(*t*) connected by *N* – 1 inextensible links of length *a* (Fig. 2), and we denote by **n***_i_*(*t*) = [**r***_i_*_+1_ (*t*) − **r*_i_***(*t*)]/*a* the unit vector pointing from bead *i* to *i* + 1. The chain is suspended in a viscous medium with shear viscosity *η* and confined within a spherical cavity of radius *R_s_*. We model the nuclear envelope, here the interior surface of the sphere, as a rigid no-slip boundary.

**FIG. 2.**
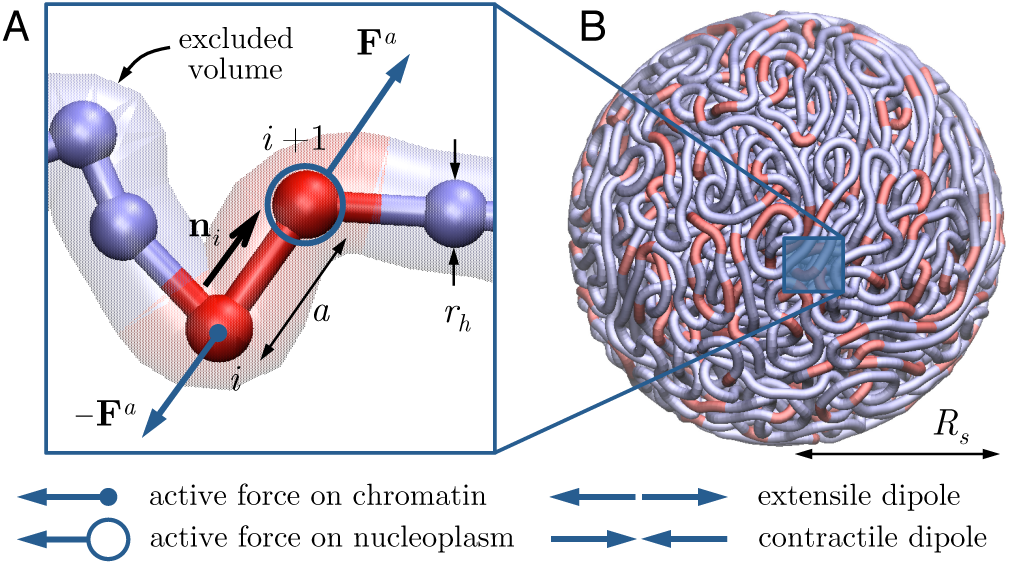
Coarse-grained chromatin model: (*A*) The chromatin fiber is modeled as a long freely-jointed chain of *N* beads each with hydrodynamic radius *r_h_* and connected by rigid links of length *a*. Active dipoles (extensile or contractile) bind stochastically to individual links, with forces applied either on the chain or on the fluid. (*B*) The chain is confined within a sphere of radius *R_s_*. In the figure, *N* = 5000 and *R_s_/a =* 8, and red segments illustrate the position of active dipoles for *P_a_ =* 1/6.

The motion of the fiber is overdamped and described by a Langevin equation for the motion of each bead [16]:

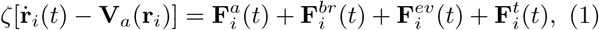

which accounts for its transport by the active nucleoplasmic flow with velocity **V***_a_*, as well as motion under the action of active forces 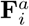, Brownian forces 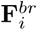, excluded volume interactions 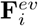, and internal tension forces 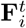. Here, *ζ* = 6*πηr_h_* is the viscous friction coefficient of one bead, equivalently expressed in terms of an effective hydrodynamic radius *r_h_*. Each chain unit in the model should be interpreted as a coarse-grained mesoscopic chromatin region consisting of a large number of nucleosomes and associated molecular motors. Brownian forces 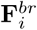 are calculated to satisfy the fluctuation-dissipation theorem, and excluded volume forces 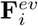 are captured by a soft repulsive potential between beads as well as links, which is truncated for |**r***_i_* − **r***_j_* | > *a* and whose strength is chosen to prevent self-crossing of the chain. Tension forces ensure that the links remain of constant length and are expressed as 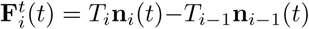, where the scalars *T_i_* are Lagrange multipliers that enforce the constraints |**n***_i_* | = 1 [17].

Fiber motion results from the action of molecular motors, which perform work on the chromatin and also drive long-ranged hydrodynamic flows. By conservation of momentum, the net active force on a chromatin segment and on the motors tethered to it must be zero, so to leading order we need only consider dipolar force pairs, which can either be extensile (←→) or contractile (→←) (Fig. 2). Active forces can either be applied directly to the chain (due to the action of the motors) or to the fluid (due to viscous drag forces on the motors themselves), or both. Here, we restrict our analysis to dipoles that are aligned with the local chain direction, and assume that one force is applied on the chain (and transmitted to the fluid via viscous drag) while the opposite force is applied directly on the fluid. The dipoles are assumed to always occur on the scale of one link, and to bind and unbind stochastically as a Poisson process with on- and off-rates *k*_on_ and *k*_off_, respectively. These two rates set the probability *p_a_* = *k*_on_/(*k*_on_ + *k*_off_) of any given link being active at an instant in time, as well as the average number *N_a_* = (*N* − 1)*k*_on_ /(*k*_on_ + *k*_off_) of active links in the system. For simplicity, we assume that all active forces have magnitude *F*_0_, corresponding to a dipole strength *F*_0_*a*. The nature of active forces in our system distinguishes it from past models for active polymers [18–21], which have typically employed isotropic colored noise to account for activity.

The active fluid flow **V***_a_* (with pressure *P_a_*) satisfies the Stokes equations for viscous hydrodynamics [22]:

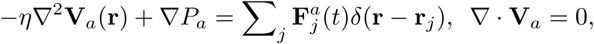

where the sum is over all active forces in the system. These equations, subject to the no-slip condition at |**r**| = *R_s_*, can be solved analytically using Green’s functions [23].

The equations of motion are integrated in time starting from a random-walk configuration for the fiber; details of the governing equations and numerical algorithm are provided in the SI. We henceforth only present dimensionless results, where variables are scaled using characteristic length scale *a* and timescale *a*^2^ζ/*k*_B_*T*, where *k*_B_*T* is the thermal energy. This is the timescale for an isolated bead to diffuse a distance of one link length by Brownian motion. With this convention, the dimensionless dipole strength is *σ* = *F*_0_*a/k*_B_*T* and compares the relative strength of activity to thermal fluctuations. We also introduce a dimensionless activity parameter

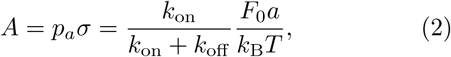

which adjusts the dipole strength for stochastic binding and unbinding with the probability *p_a_* of a link being active.

## RESULTS AND DISCUSSION

### Unconfined active chains

To gain intuition, we first consider the time evolution of short unconfined chains. Figure S1 of the SI compares a Brownian passive chain to two active chains, one with tethered contractile dipoles and the other with extensile dipoles; also see SI movies. Their differences are plain. The contractile case is similar to the Brownian case albeit with enhanced fluctuations induced by activity; chain conformations remain random and disorganized. The extensile case, however, is characterized by the gradual straightening of the chain through hydrodynamic and tensile interactions with itcelf, which in competition with thermal fluctuations results in a persistence length well beyond the chain length. During the transient, the simulation shows persistent folded states that arise due to nonlocal extensile flows leading to alignment of nearby stretches of fiber. These states slowly unbend at the folds as the chain progressively unravels.

### Chromatin model: Confined chains

We now turn to the dynamics of longer chains under confinement. All simulations shown here are for a fiber composed of *N* = 5000 beads in a confining sphere of radius *R_s_/a* = 8.0; also see SI movies. To set a baseline, Fig. 3A (*top row*) shows the dynamics of a single fluctuating, spherically confined fiber in the absence of active motors. During a brief initial transient, steric forces push nearly overlapping regions of the fiber away from each other, and steric and fluctuation forces come roughly into balance leading to a roughly uniform density (see Fig. S2). The macroscopic dynamics in this phase is extremely slow: examination of the frames shows the chain as essentially frozen in place, as its motion is strongly hindered by its self-entanglement and confinement. While excluded-volume interactions do lead to some local nematic alignment of spatially proximal stretches of fiber (Fig. 3B, *top row*), it is spatially disorganized as is the fiber displacement field (Fig. 3C, *top row*). When instead contractile dipoles are active (Fig. 3, *middle row*), the chain now shows more significant displacements (Fig. 3A) but again little long-range ordering is evident (Fig. *3B*), though there is more coherence in displacement fields on intermediate length scales than in the Brownian case (Fig. 3*C*).

**FIG. 3.**
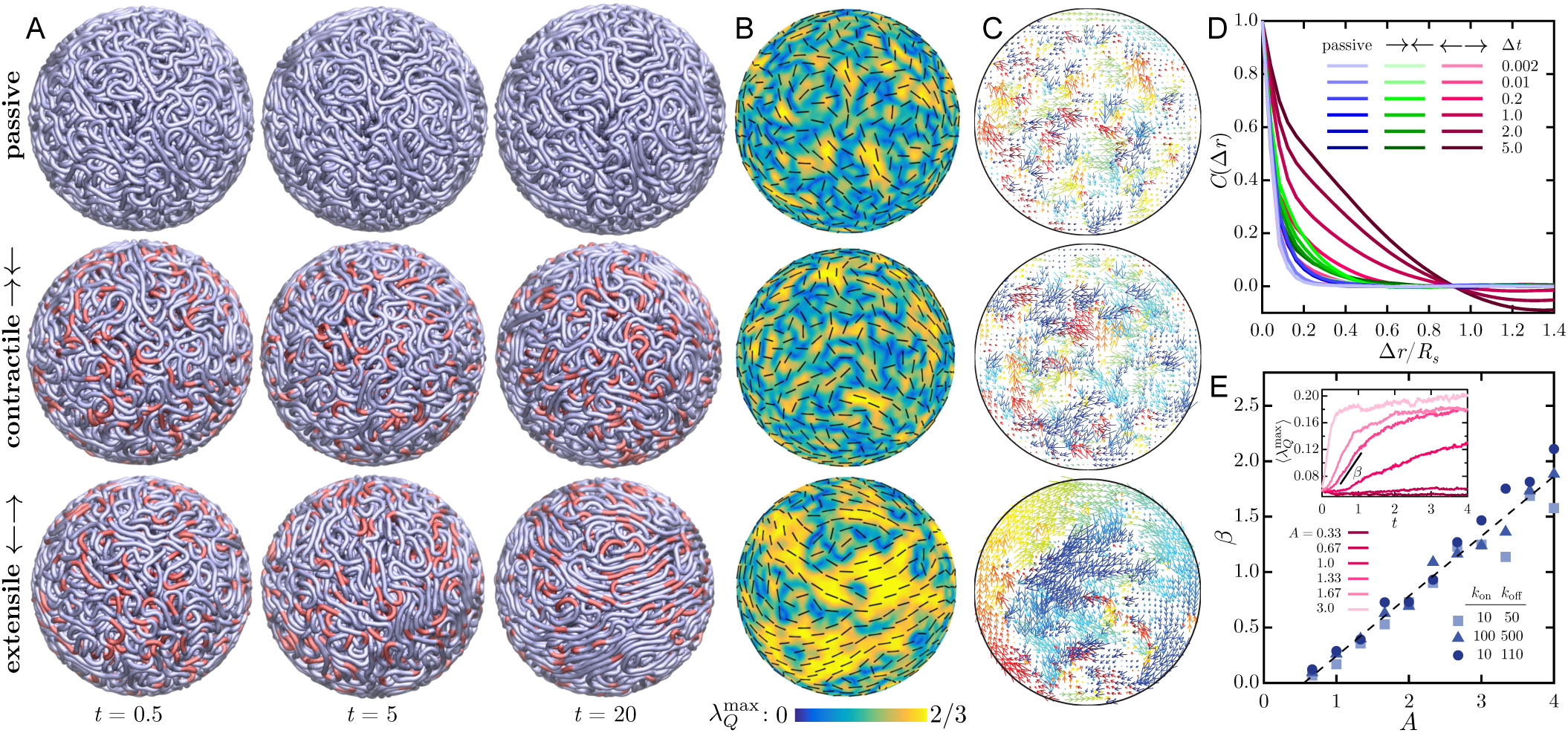
Simulations of a passive Brownian chain (*top row*) and active chains under the action of contractile (→←, *middle row*) and extensile (←→, *bottom row*) force dipoles, with *N* = 5000 and *R_s_/a* = 8. In the, active cases, *A* = 2, *k*_on_ = 10, *k*_off_ = 50. Also see SI movies. (*A*) Fiber configurations at different times with red segments showing instantaneous positions of active dipoles. (*B*) The nematic structure of the fiber in a shell of thickness *a* on the surface of the confining sphere. Colors show the maximum eigenvalue 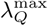 of the local nematic order tensor **Q**(**r**) = 〈**nn** – **I**/3〉 (i.e., the scalar order parameter), and the black segments show the corresponding nematic director. (*C*) Chromatin displacement maps calculated in a plane across the spherical domain over a time interval Δ*t* = 0.2. Arrows are color-coded according to the direction of displacement. (*D*) Average spatial autocorrelation functions of the displacement maps for increasing values of the displacement time Δ*t*. (*E*) Growth rate *β* of the order parameter 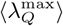 as a function of activity *A* for different combinations *k*_on_ acd *k*_off_. The growth rate is calculated during the initial exponential increase of the order parameter, which we average over spherical regions of size 0.5*R_s_* (see inset).

The dynamics is very different when extensile motors are active (Fig. 3*A, bottom row*). As in the other cases, there is first a brief rearrangement period during which the initially dominant steric repulsion forces are relaxed. Following this, the system rapidly reorganizes itself as the chromatin fiber draws itself out into long, mutually aligned segments. This alignment has its origin in motor activity: long-ranged nucleoplasmic flows are generated along the chain by motor activity and tend to both straighten the fiber locally and draw nearby regions into alignment. In a fashion reminiscent of other extensile active systems [5, 6], these large-scale reconfigurations involve a positive feedback loop by which the cooperative alignment of elongated active chain segments is reinforced by the hydrodynamic flows they generate. The result is the appearance of large regions with high nematic order in fiber orientation (Fig. 3B, *bottom row*), which coarsen with time as the chain continues to unfurl and which are separated by sparse disorganized regions. The dynamics near the boundary in this case bears resemblance to that of active 2D nematics on spherical surfaces [24–26], yet is fundamentally different due to the strong hydrodynamic and mechanical couplings that exist with the nucleus interior.

These internal dynamics and rearrangements have clear signatures in the displacement maps of Fig. 3*C*. While passive Brownian chains exhibit random uncorrelated displacements similar to those seen in ATP-depleted nuclei (Fig. 1*B*, *bottom row*), short-scale motions are visible in contractile systems due the local flows induced by individual dipoles. Extensile systems, however, exhibit coherent chain motions on large length and time scales resembling those in wildtype nuclei (Fig. 1*B*, *top row*). We note that motor-induced nucleoplasmic flows alone are sufficient to qualitatively produce these dynamics. These flows have their origin, again, in the induced alignment of fiber segments by extensile motor activity: by aligning the fiber across the system, more and more motors are recruited into producing aligning flows. That is, large-scale fluid and material flows and fiber alignment are tightly coupled and self-reinforcing aspects of the system dynamics. This is especially reminiscent of the dynamics observed in active particle suspensions where extensile stresses are predominant and alignment forces are strong [7, 13].

These observations are made quantitative in Fig. 3*D*, showing in each case the average spatial autocorrelation functions of displacement maps for different time intervals Δ*t*. The Brownian chain shows no correlation on length scales greater than one link length, and is reminiscent of ATP-depleted nuclei (Fig. 1*C*, *blue lines*). Contractile active chains do exhibit correlations on short length scales of the order of a few link lengths, which are strongest at an intermediate time scale of Δ*t* ≈ 0.2 corresponding approximately to the duration of individual active events. In extensile systems, correlations occur on the system scale and persist for very long times in agreement with the dynamics observed above, and are similar to those measured in wildtype nuclei in Fig. 1*C* (*red lines*).

The temporal growth of large-scale conformational changes is characterized in Fig. 3*E*. The mean scalar nematic order parameter 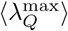, defined as the maximum eigenvalue of the nematic order tensor **Q(r)** = 〈**nn** − **I**/3〉 averaged over spherical subdomains of diameter 0.5*R_s_*, is plotted in the inset and exhibits an initial exponential growth followed by saturation. The corresponding growth rate *β* in rapid growth regime, plotted vs *A* in the main figure, shows a continuous transition to coherent motion above a finite level of activity. A detailed mapping of the (*p_a_,σ*) space in Fig. S3 confirms the existence of a well-defined transition that is governed purely by the net activity of the system and occurs at *A_c_* ≈ 0.4. This transition likely signals a supercritical instability of the initial random state akin to that of extensile active suspensions [14, 15]. In unstable systems, the initial growth rate increases linearly with *A* and is largely unaffected by dipolar on/off rates.

The nature of nematic alignment at late times in the simulations is detailed in Fig. 4*A*, showing the scalar order parameter averaged over spherical domains of radius r. Both passive and contractile systems show high nematic order only on the scale of one link (*a/R_s_* = 0.125) as a result of excluded volume. On the other hand, all the extensile cases show much extended order on the system scale, as would be inferred from Fig. 3. This difference is reflected in the spatial autocorrelation function of the director **n***_i_* along the chain (Fig. 4*B*), which for the extensile cases shows anti-correlation in orientation once the chain has spanned the radius of the sphere (which is 8*a* across). This anti-correlation is a direct consequence of confinement, which frustrates the ability of the chain to stretch on length scales greater than the system size and constrains it to curve around; this qualitative picture is further clarified in Fig. 5.

**FIG. 4.**
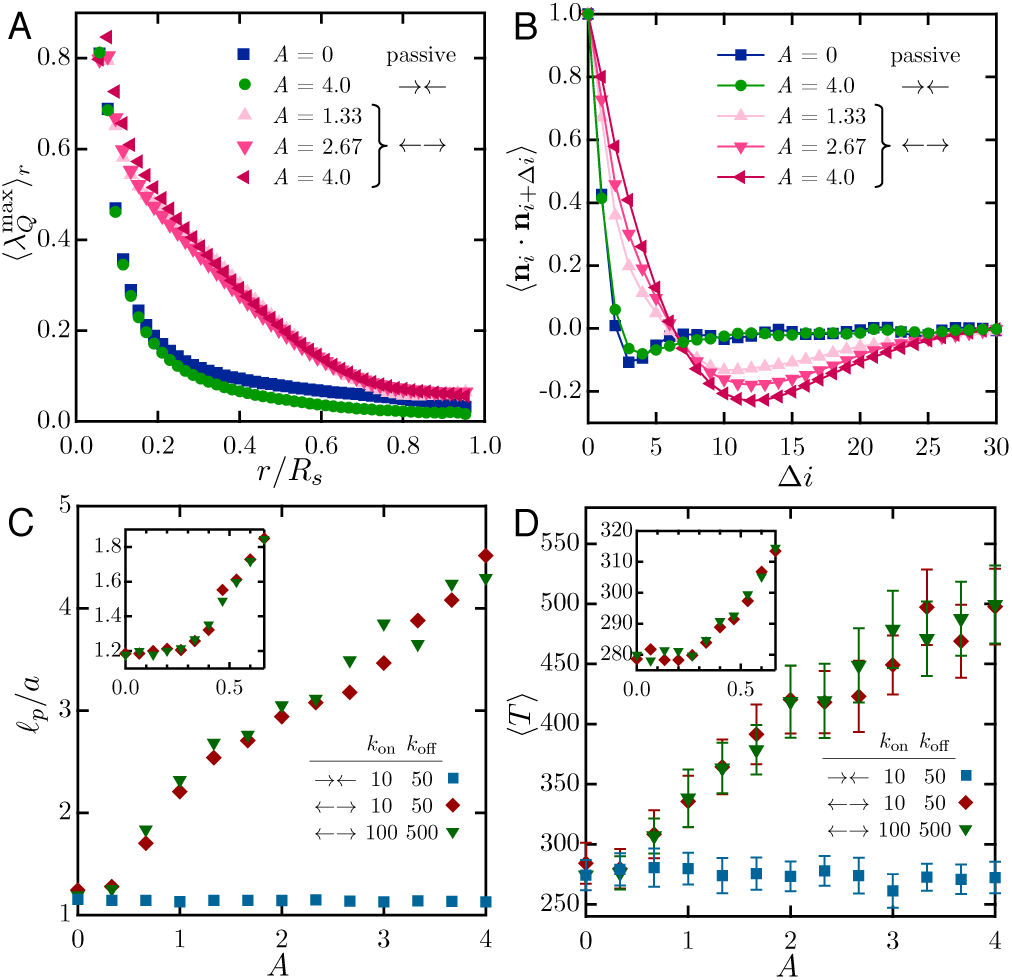
(*A*) The scalar nematic order parameter 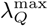, averaged over spheres of radius *r*, at late times in simulations of passive, contractile (→←) and extensile (→←) systems. (*B*) Autocorrelation function of the link orientation vector *n* (where *i* is the link index) as a function of separation distance Δ*i* along the fiber. (*C*) Effective persistence length *ℓ_p_* as a function of activity in contractile and extensile systems. Inset highlights the birfurcation occurring near *A* ≈ 0.4 in excellent agreement with the phase diagram of Fig. S3. (*D*) Mean scalar tensile force 〈*T*〉 inside the chain as a function of A in the same systems.

**FIG. 5.**
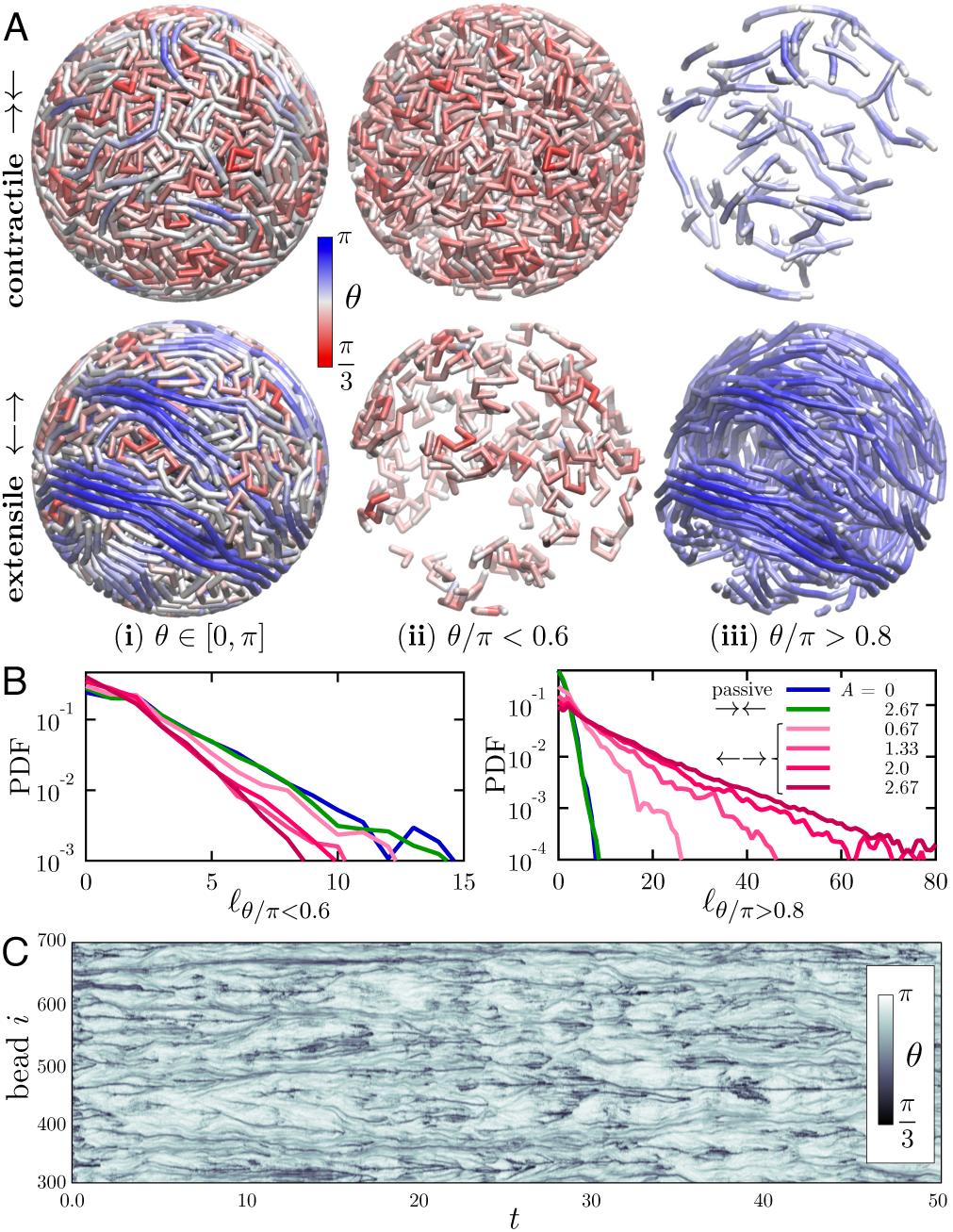
(*A*) Snapshots of contractile and extensile systems (*A* = 1) where the chain is colored according to the local bond angle *θ* = π − cos^−1^ (**n***_i_* · **n***_i_*_+1_). The entire chains are shown in (**i**), whereas (**ii**) and (**iii**) isolate high- and low-curvature chain segments, defined as portions of the chains where *θ*/π < 0.6 and > 0.8, respectively. (*B*) Probability density functions of the length of high- and low-curvature segments for passive, contractile and extensile systems. (*C*) Spatiotemporal diagram showing the evolution of the bond angle distribution along chain subset *i* ∈ [300, 700] in an extensile system with *A* = 1.

In addition to driving coherent flows and displacements, motor activity also modifies the effective mechanical properties of the fiber. First, by straightening local chain segments, extensile activity confers an effective bending rigidity to the chain. We quantify this effect in Fig. 4*C*, where we calculate the chain persistence length *ℓ_p_/a* = (1+ξ)/2(1−ξ) with ξ = (**n***_i_*·**n***_i_*_+1_) [27]. While *ℓ_p_/a* remains close to 1 for all levels of activity in contractile systems, extensile chains show a linear increase in persistence length with *A* and thus in effective bending rigidity above the transition to coherent motion. Internal tensile forces are also affected (Fig. 4*D*), with a mean tension 〈*T*〉 that also continuously increases from its baseline in extensile systems.

The internal structure and conformation of these confined chains can be gleaned in more detail by analyzing the spatial distribution of chain curvature or, equivalently, bond angle *θ_i_* = *π −* cos^−1^(**n***_i_* · **n***_i_*_+1_) in Fig. 5*A*. In contractile systems, the chain is primarily bent with small bond angles (*red*) throughout the system, with the occasional occurrence of short straight segments (*blue*). The structure is quite different in extensile systems: in that case, regions of high curvature are sparse and segregated, while highly organized extended chain segments fill large spatial subdomains inside the sphere. The distribution of lengths of highly-bent segments (*θ/π <* 0.6) in Fig. 5*B* depends only weakly on activity and decays very rapidly, with most segments only involving a few links. Extensile activity, however, strongly increases the probability of long straightened segments (*θ/π >* 0.8). Their length is found to follow an exponential distribution, indicating Poissonian statistics for the occurrence of kinks and entanglements along the chain. Extended segments with lengths up to 160 links are observed in some simulations, implying a highly organized spatial structure in which long chain portions straightened by activity-driven flows wrap around and across the system as anticipated from Fig. 4*B*. The dynamic nature of fiber curvature in the extensile case is highlighted in Fig. 5*C*, showing the spatiotemporal evolution of the bond angle distribution along a section of 400 beads. Most initial kinks and turns present at *t* = 0 quickly disappear as chain segments start to stretch and straighten under self-induced flows. Remaining highly bent regions (*black* streaks in Fig. *5C*) dynamically merge, annihilate and nucleate, though they remain mostly localized along the chain. Several of them are seen to persist on very long time scales and correspond to frustrated topological entanglements. In extreme cases, these entanglements come in the form of pseudo-knots that were present in the initial chain configuration and are made tighter by extensile activity.

## CONCLUSIONS

The essence of our chromatin model is its mechanical coupling of a very flexible extended polymer in a confined fluidic environment moving under dipolar forces created by the activity of tethered motors. It is when the resulting active dipoles are extensile that large-scale coherent motion is observed, and the emergence of this coherence is intimately related to non-local chain stretching and alignment driven by hydrodynamics. While our model suggests that hydrodynamic interactions between dipolar forces alone can lead to the large-scale coherent motion observed *in vivo*, it is conceivable that electrostatic as well as van der Waals interactions might contribute to the effective amplitude of dipolar forces at short length scales [28–31]. In addition to nonspecific physical interactions, proteins involved in chromatin remodeling (e.g. SWI/SNF, ISWI) and structural maintenance of chromosomes (e.g. cohesin) could also be affecting the length and time scale of the coherent motions by generating bends, folds, entanglements or loops in the chromatin fiber [32–35].

The precise biological function of the coherent chromatin motion remains to be revealed, and, in principle, could simply be a side effect of other dynamic biological processes inside the nucleus. Still, these dynamics involve the collective motion of genes on time scales of seconds, and it seems more likely that these displacements have major implications for the spatiotemporal organization of the genome. For example, the organized nucleoplasmic flows generated by the dipolar forces could contribute to gene regulation by facilitating the distribution of the transcription machinery in the cell nucleus through advective instead of diffusive transport. To connect the extensile activity with its biological origins, we need to continue building and bridging microscopic and macroscopic understandings of chromatin dynamics, both experimentally and theoretically.

## MATERIALS AND METHODS

### Cell Culture and Biochemical Perturbations

Stable HeLa H2B-GFP cell line (CCL-2) was cultured according to American Type Culture Collection (ATCC) recommendations. Cells were cultured in a humidified, 5% CO_2_ (vol/vol) atmosphere at 37^o^ C in Gibco Dulbecco’s modified eagle medium (DMEM) supplemented with 10% FBS (vol/vol), 100units/mL penicillin, and 100mg/mL streptomycin (Invitrogen). Before the experiment cells were plated on 35-mm Mat-Tek dishes with glass bottom No. 1.5 (MatTek) for 24 h and the medium was replaced by Gibco CO_2_-independent medium supplemented with L-glutamine (Invitrogen). Cells were then mounted on the microscope stage kept in a custom-built 37^o^C microscope incubator enclosure with 5% CO2 (vol/vol) delivery during the entire experiment. To deplete ATP, cells were treated with 6mM 2-deoxyglucose (Sigma Aldrich) and 1 *μ*M trifluoromethoxy-carbonylcyanide phenylhydrazone (Sigma Aldrich) dissolved in CO_2_-independent medium supplemented with L-glutamine 2 h before imaging.

### Microscopy, Image Acquisition and DCS

Images were acquired and DCS analysis was carried out following procedures previously described in [1]. For detailed microscopy and image acquisition protocol see *Supporting Information*. DCS maps and *C*(Δ*r*) were calculated for Δ*t* = 0.25 s −15 s for 10 wildtype and 10 ATP-depleted nuclei following procedures from [1].

## ACKNOWLEDGMENTS

DS acknowledges support from National Science Foundation (NSF) Grant DMS-1463965. MJS acknowledges support from NSF Grants DMS-1463962, DMS-1620331, as well as the NYU MRSEC Grant DMR-1420073. AZ is grateful for the support from the National Institutes of Health Grant R00-GM104152, NSF CAREER Grant PHY-1554880 and Whitehead Fellowship for Junior Faculty in Biomedical and Biological Sciences.

